# CFTR function in alveolar type 1 cells drives lung liquid secretion and host defense

**DOI:** 10.1101/2025.03.25.645303

**Authors:** Sayahi Suthakaran, Sonya Homami, Deebly Chavez, Stephanie Tang, Sarah K.L. Moore, Chaya Sussman, Jimmy Zhang, Clemente J. Britto, Alice Prince, Alison J. May, Jaymin J. Kathiriya, Jaime L. Hook

**Author notes:** These authors contributed equally to this work (Sayahi Suthakaran, Sonya Homami).

## Abstract

Loss of the liquid layer that lines the lung’s air-facing surface underpins mechanisms of major lung diseases, but the development of therapies that restore liquid secretion is hampered by an incomplete understanding of the cell types that drive it. Here, we show CFTR function in alveolar type 1 (AT1) cells – a cell type that comprises 95% of the lung surface but is presumed to be unimportant in CFTR-related diseases – is critical to lung liquid secretion and the secretion-mediated clearance of particles and *S. aureus* from lung alveoli. Our findings reveal essential roles for AT1 cells in lung homeostasis and defense, and they call for a reevaluation of the role of AT1 cells in CFTR-related diseases. We suggest AT1 cells be considered key targets of secretion-restoring therapies.

## Main Text

Loss of the liquid layer that lines the lung’s epithelial surface contributes to the pathogenesis of cystic fibrosis (CF) (*1, 2*), chronic obstructive pulmonary disease (COPD) (*3, 4*), and lung infection (*5–7*). Therapeutic approaches to rescue liquid secretion are designed to act in the lung airways (*8–10*), where liquid secretion has been traditionally identified. However, lung alveoli comprise over 95% of the lung’s air-facing surface area and continuously secrete liquid (*6, 11–13*), indicating alveoli might be an important source of lung liquid secretion and an unrecognized therapeutic target. The development of alveolus-targeted therapies has been hampered by a lack of understanding of the cellular source of alveolar liquid secretion and its relevance to lung health.

Alveolar liquid secretion depends on function of the cystic fibrosis transmembrane conductance regulator (CFTR) protein in the alveolar epithelium (*6, 7, 13–15*). Alveolar type 2 (AT2) cells express CFTR (*14–16*) and are thought to drive alveolar liquid dynamics. However, alveolar type 1 (AT1) cells also express CFTR and comprise nearly all of the alveolar surface (*17–20*), indicating their contribution to alveolar liquid secretion might be considerable. Yet, there is little understanding of the contribution of AT1 cells to alveolar liquid secretion, in part because their low *Cftr* mRNA content (*17*) suggests it is negligible.

We quantified cell-specific contributions of CFTR function to alveolar liquid secretion using confocal microscopy of live, intact, perfused lungs of transgenic mice with AT1- and AT2-specific expression of a *Cftr* null allele. Our findings show AT1 cell CFTR function drives homeostatic alveolar liquid secretion and the secretion-mediated clearance of particles and bacteria from alveoli. These data provide the first evidence that AT1 cells are essential to lung liquid secretion and defense and should be considered critical targets of secretion-rescuing therapies. Moreover, our findings call for a reevaluation of the role of AT1 cells in CFTR-related diseases traditionally thought to center on the airways.

### AT1 and AT2 cell-specific transgene expression

We used established transgenic mouse strains to generate mice in which CFTR^null^ and GFP expression were tamoxifen-induced in adulthood in an AT1 or AT2 cell-specific manner. AT2-CFTR^null^-GFP mice carried *Sftpc-CreER^T2^*(*21*), *Cftr^fl10/fl10^* (*22*), and *ROSA^mT/mG^*, a reporter allele that confers membrane-targeted TdTomato expression prior to Cre-mediated excision and GFP expression after (*23*). Likewise, AT1-CFTR^null^-GFP mice carried *Ager-CreER^T2^* (*24*), *Cftr^fl10/fl10^*, and *ROSA^mT/mG^*; AT2-AT1-CFTR^null^-GFP mice carried *Sftpc-CreER^T2^, Ager-CreER^T2^*, *Cftr^fl10/fl10^*, and *ROSA^mT/mG^*; and littermate control (LMC-TdTomato) mice carried *Cftr^fl10/fl10^* and *ROSA^mT/mG^*.

We used our standard methods (*7, 25*) to view live alveoli of intact, perfused lungs of AT2-CFTR^null^-GFP mice by real-time confocal microscopy. GFP fluorescence was evident in plasma membranes of rounded cells located primarily at septal junctions (**Fig. 1**), consistent with AT2 cell morphology and location (*26, 27*). To test AT2 cell function, we instilled alveolar airspaces with LysoTracker dye, which localizes to AT2 cell lamellar bodies and signals inflation-induced surfactant secretion (*28*). Lung hyperinflation caused time-dependent loss of LysoTracker fluorescence in GFP-positive cells (**fig. S1, A-B**), indicating GFP-positive cells function as AT2 cells. To quantify *Cftr* deletion efficiency, we isolated AT2 cells (**fig. S2, A-C**) and performed qPCR of *Cftr* RNA. AT2 cells of an AT2-CFTR^null^-GFP mouse had 100-fold reduction of *Cftr* RNA compared with AT2 cells of an LMC-TdTomato mouse (**Table S1**). Taking these findings together, we conclude that AT2 cells of AT2-CFTR^null^-GFP mice are *Cftr*-deficient and marked by GFP fluorescence.

**Fig. 1.**
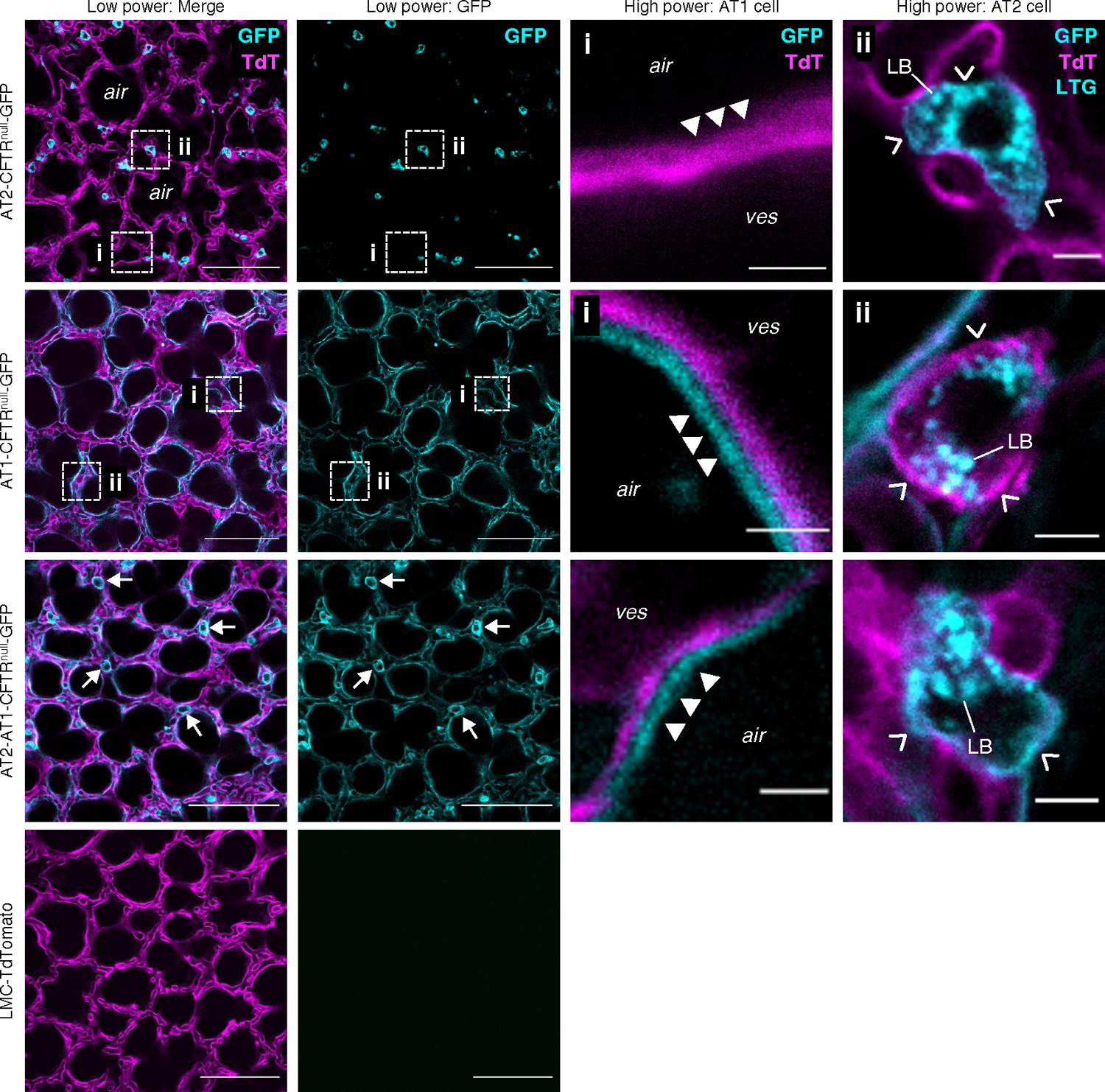
Alveolar epithelial cell type-specificity of GFP expression. Confocal images show plasma membrane *GFP* and TdTomato (*TdT*) fluorescence in live alveoli of intact, perfused lungs of mice of the indicated genotype. *Dashed squares* show locations of the high power views shown in the third and fourth columns. *Closed arrowheads* point out AT1 cell fluorescence, which is GFP+ (*cyan*) and distinct from neighboring TdT+ (*magenta*) endothelial cells only in AT1-CFTR^null^-GFP and AT2-AT1-CFTR^null^-GFP mice. *Open arrowheads* show fluorescence of AT2 cells, which is GFP+ (*cyan*) only in AT2-CFTR^null^-GFP and AT2-AT1-CFTR^null^-GFP mice. In the fourth column, cytosolic fluorescence of lysotracker green (*LTG*) delineates lamellar bodies (*LB*) and confirms AT2 cell identity. In the third row, *arrows* show example AT2 cells in an AT2-AT1-CFTR^null^-GFP mouse. *Air,* airspace; *ves,* microvessel lumen. Scale bars: 100 (*first* and *second columns*), 3 (*third columns*), and 5 (*fourth columns*) µm. Images were replicated in lungs of 3 mice per genotype. Note, in the fourth column, both LTG and GFP are pseudocolored cyan.

By contrast, GFP fluorescence in live lungs of AT1-CFTR^null^-GFP mice was evident in thin cells that line alveolar walls (**Fig. 1**), consistent with AT1 cell morphology and location. AT2 cells had abundant *Cftr* RNA (**fig. S2D and Table S1**). Taking these data together, we interpret that CFTR^null^ expression in AT1-CFTR^null^-GFP mice is AT1 cell-specific.

In AT2-AT1-CFTR^null^-GFP mice, GFP fluorescence was evident in plasma membranes of both AT2 and AT1 cells (**Fig. 1**), and AT2 cell qPCR showed more than 100-fold reduction of *Cftr* RNA (**fig. S2E, Table S1**). There was no GFP fluorescence in alveoli of LMC-TdTomato mice (**Fig. 1**). We interpret that AT2-AT1-CFTR^null^-GFP mice are *Cftr-*deficient in both AT2 and AT1 cells, while LMC-TdTomato mice are *Cftr*-deficient in neither cell type.

### CFTR function in AT1 cells drives alveolar liquid secretion

We imaged alveolar wall liquid (AWL) secretion in real-time in live lungs by microinstilling alveolar airspaces with fluorophore-tagged dextran (*6, 7*). Time-dependent loss of dextran fluorescence signals dextran dilution by secreted AWL (*6, 7*). As expected, microinstilled dextran pooled at structural niches of alveoli, where septa converge (**Fig. 2A**). Also as expected, dextran fluorescence decreased in a time-dependent manner in alveoli of LMC-TdTomato mice (**Fig. 2, B-C**), signaling the alveolar epithelium secreted AWL. To rule out the possibility that the fluorescence loss resulted from dilution by airspace edema fluid, we added fluorophore-tagged dextran to the lung perfusate solution (*7, 25*). Perfusate fluorescence localized to microvessels but not airspaces (**Fig. 2D**), indicating alveolar barrier function was intact and ruling out edema as the cause of the fluorescence loss. We conclude that AWL secretion caused time-dependent loss of dextran fluorescence in alveoli of LMC-TdTomato mice.

**Fig. 2.**
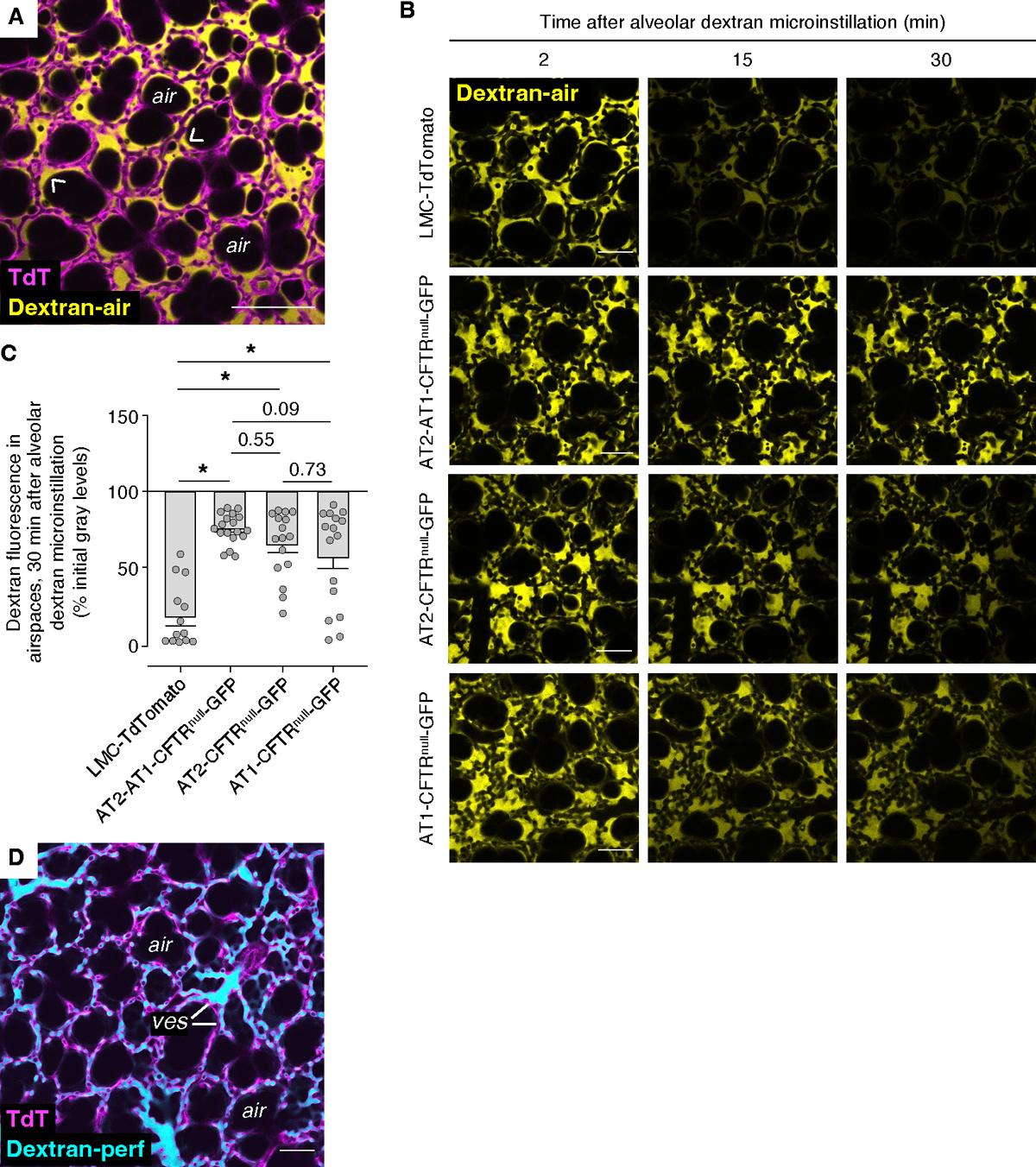
Real-time confocal imaging of liquid secretion in live alveoli. (**A**) Images show fluorescence of AlexaFluor 647-conjugated dextran (*Dextran-air*; 10 kD; 10 mg/mL; *yellow*) in airspaces of live, perfused alveoli (*magenta*) of an LMC-TdTomato mouse at 5 min after alveolar dextran microinstillation. Note, dextran formed a thin layer against alveolar walls and pooled at niches (examples: *arrowheads*). *TdT*, TdTomato; *air,* example airspace. Scale bars: 100 µm. Replicated in 12 mice. (**B-C**) Images (B) and group data (C) show time-dependent change of dextran fluorescence in airspaces of live alveoli in lungs of mice of the indicated genotypes. Fluorescence of alveolar walls is not shown. In C, circles each represent mean dextran fluorescence in one imaging field of at least 30 alveoli; bars: mean ± SEM; *n*=3 mice per group; *p* as indicated or < 0.05 (*) by ANOVA with post hoc Tukey testing. Scale bars: 50 µm. (**D**) Images show fluorescence of TdTomato (*TdT; magenta*) in alveoli and fluorescein isothiocyanate (FITC)-labeled dextran (*Dextran-perf*; 20 kD; 5 mg/mL; *cyan*) in the perfusate solution of live alveoli of intact, perfused lungs of an LMC-TdTomato mouse. Scale bar: 50 µm. Replicated in 12 mice.

By contrast, dextran fluorescence loss was blocked in alveoli of AT2-AT1-CFTR^null^-GFP mice (**Fig. 2, B-C**), affirming that AWL secretion results from CFTR function in the alveolar epithelium (*6, 7*). While airspace dextran fluorescence also decreased in alveoli of AT2-CFTR^null^-GFP and AT1-CFTR^null^-GFP mice, the rates of fluorescence loss were majorly reduced compared with LMC-TdTomato mice (**Fig. 2, B-C**). Dextran perfusion studies showed normal barrier function in all mice (data not shown). These findings show *Cftr* deficiency in each of AT2 and AT1 cells partially blocked AWL secretion, indicating that CFTR function in both AT2 and AT1 cells generates AWL.

Heterogeneous recombination efficiency across alveoli of tamoxifen-treated *Ager-CreER^T2^* mice (*24*) provides an opportunity to relate the magnitude of AT1 cell CFTR^null^ expression to AWL secretion rates. In lungs of AT1-CFTR^null^-GFP mice, we imaged alveolar regions with low, medium, or high AT1 cell GFP fluorescence (**Fig. 3, A-B**), which we interpreted to indicate low, medium, and high CFTR^null^ expression, respectively. In low-GFP regions, airspace dextran fluorescence decreased rapidly (**Fig. 3C**). Fluorescence loss was slower in medium-GFP regions (**Fig. 3C**) and slower still in high-GFP regions (**Fig. 3C**). These findings indicate the extent of AT1 cell GFP fluorescence relates directly to AWL secretion rates (**Fig. 3D**), such that alveolar regions with the most CFTR^null^ expression generated the least secretion, and vice versa. This direct, “dose-dependent” relationship between AT1 cell CFTR expression and AWL secretion strengthens our conclusion that AT1 cell CFTR function is a major driver of alveolar liquid secretion.

**Fig. 3.**
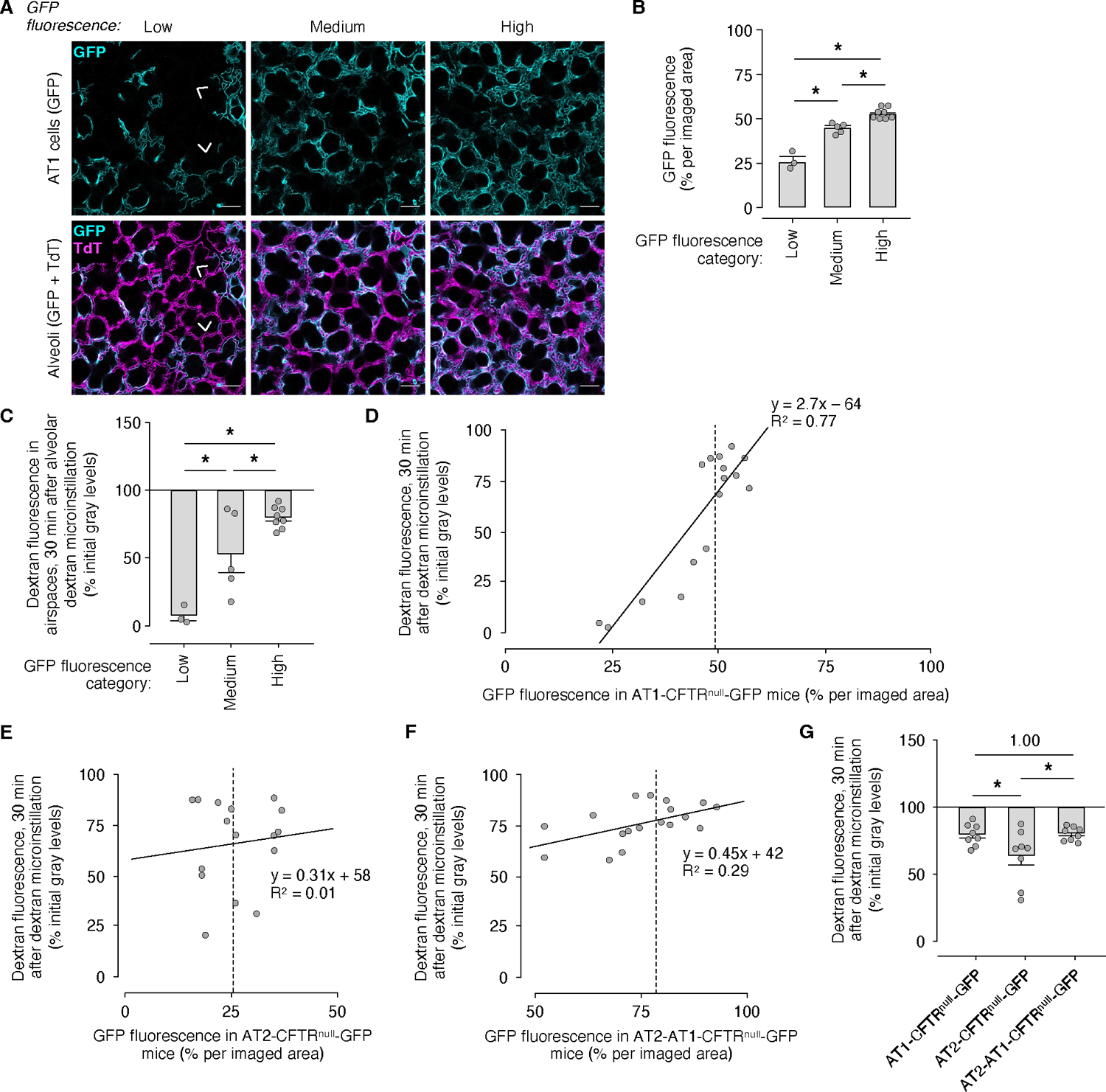
“Dose-dependent” relationship between AT1 cell *Cftr* deficiency and inhibition of alveolar liquid secretion. (**A-B**) Confocal images (A) and group data (B) show heterogeneity of *GFP* fluorescence across imaging fields of at least 30 live alveoli in intact, perfused lungs of AT1-CFTR^null^-GFP mice. TdTomato (*TdT*) fluorescence in GFP-free alveoli (examples: *arrowheads*) shows some AT1 cells expressed TdTomato, but not GFP. In B, circles each represent mean GFP fluorescence in one imaging field; bars: mean ± SEM; \**p* < 0.05 by ANOVA with post hoc Tukey testing. Each group includes imaging fields generated in at least 2 mice. Scale bars: 50 µm. (**C-G**) Group data quantify change of AlexaFluor 647-conjugated dextran fluorescence after alveolar dextran microinstillation into airspaces of the imaging fields shown in panels A and B (C-D) or into airspaces of mice of the indicated genotype (E-G). Circles each represent mean dextran fluorescence in one imaging field; N=3 mice per genotype; bars: mean ± SEM; \**p* < 0.05 by ANOVA with post hoc Tukey testing. In D-F, *solid lines* were calculated by linear regression, and *dashed lines* set apart the 8 imaging fields with high GFP fluorescence.

We followed up by comparing dextran fluorescence loss in high-GFP regions in lungs of AT1-CFTR^null^-GFP, AT2-CFTR^null^-GFP, and AT2-AT1-CFTR^null^-GFP mice (**Fig. 3, D-F**). Our findings show inhibition of dextran fluorescence loss was equivalent in AT1-CFTR^null^-GFP and AT2-AT1-CFTR^null^-GFP mice, but less in AT2-CFTR^null^-GFP mice (**Fig. 3G**). Because the maximum inhibition of AWL secretion achievable with AT1 cell CFTR^null^ expression (**Fig. 3, D and G**) exceeds that with AT2 cell CFTR^null^ expression (**Fig. 3, E and G**), we conclude that AT1 cells are the primary source of AWL.

### AT1 cell CFTR function clears particles from alveoli

AWL secretion defends against particle stabilization in alveoli by convectively transporting particles toward the airways (*6*). We considered AT1 cell AWL secretion might be essential to particle clearance mechanisms. To test this possibility, we used an established method (*6, 25*) to microinstill fluorescent polystyrene beads (1 μm diameter) into airspaces of medium-GFP regions of AT1-CFTR^null^-GFP mice. We analyzed the initial organization and movement of 80 bead groups by generating confocal images of the beads from 30 min to 3 h after microinstillation, then measuring their displacement according to: (A) their initial location, defined as bead position against a flat alveolar septum or a niche, a corner-like structure where septa converge; and (B) their initial group size, defined as a small cluster (1-3 beads) or a microaggregate (4 or more beads).

Our findings show the beads formed small clusters and microaggregates at flat septa and niches within minutes of microinstillation (**Fig. 4A**). Across imaged regions of alveoli, there was no difference in the number of bead groups that initially localized to alveolar walls lined by TdTomato- or GFP-expressing AT1 cells (**Fig. 4B**). The distribution of bead subgroups – i.e., small clusters at flat septa, microaggregates at flat septa, small clusters at niches, and microaggregates at niches – was also equivalent (**fig. S3A**). However, mean group size at TdTomato-expressing AT1 cells was greater (**Fig. 4C**), because microaggregates at TdTomato-expressing AT1 cells were larger (**fig. S3B**) and incorporated more beads. We interpret that AT1 cell CFTR^null^ expression did not influence how microinstilled beads initially organized against alveolar walls, but it might have had an inhibitory effect on bead microaggregation.

**Fig. 4.**
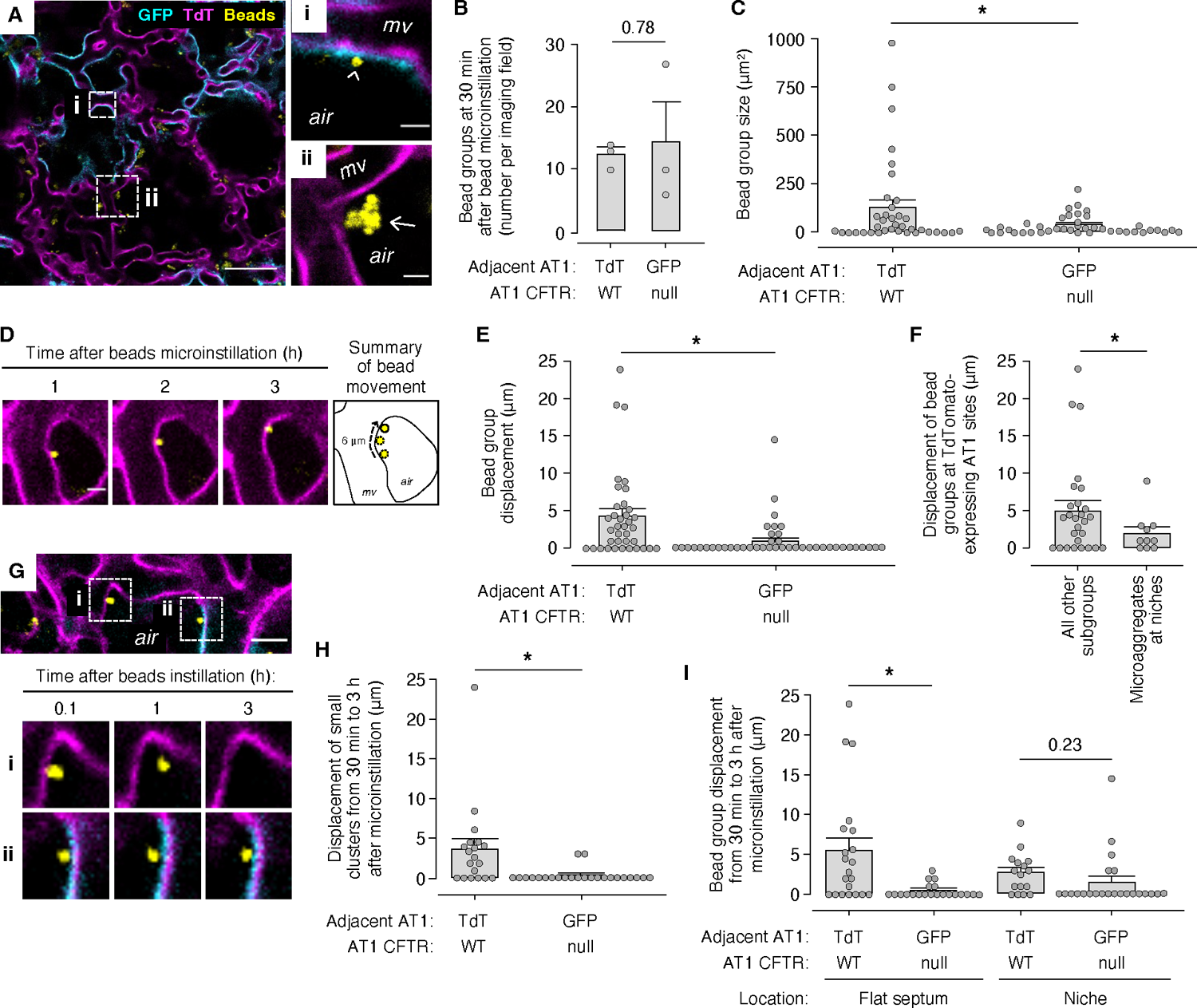
AT1 cell CFTR function drives particle clearance in alveoli. We microinstilled airspaces of live, perfused alveoli of AT1-CFTR^null^-GFP mice with fluorescent beads (*yellow*). **(A-C)** Confocal images (A) and group data indicate initial locations (B) and sizes (C) of the bead groups at 30 min post-instillation. Bead groups were adjacent to AT1 cells that expressed GFP (*cyan*) or TdTomato (*TdT; magenta*), and text in the group data indicates CFTR expression (CFTR^null^ or WT) signaled by GFP or TdT fluorescence. *Dashed squares* show locations of the high-power views shown in *i* and *ii*. *Arrowhead* (i) and *arrow* (ii) show, respectively, a small cluster adjacent to a GFP-expressing flat septum and a microaggregate at a TdTomato-expressing alveolar niche. *air*, airspace; *mv*, microvessel. (**D-I**) Follow-up images (D,G) and group data (E,F,H,I) quantify convective displacement of beads along alveolar walls after microinstillation. In the group data (B-C, E-F, H-I), circles indicate *n* and each represent the number of bead groups in one imaging field of at least 30 alveoli (B) or single bead groups (all others); bars: mean ± SEM; *p* as indicated or < 0.05 (*) by two-tailed *t* test. Scale bars: 30 (A), 3 (i, ii, D), and 15 (G) µm.

We next analyzed the movement of beads located at TdTomato-expressing AT1 cells (**Fig 4D**). Our data show the beads were spontaneously displaced a mean distance of 4 µm along alveolar walls during the imaging period (**Fig. 4, D-E**), confirming that beads are convectively transported in alveoli (*6*). The shortest displacement distances occurred among microaggregates at niches (**Fig. 4F**), affirming our published findings (*25*) that niche microaggregates resist alveolar clearance mechanisms.

Although most bead groups displaced from flat surfaces were transported toward the airways, one-third moved toward niches (data not shown). At the same time, niche microaggregates tended to enlarge (**fig. S3, C-D**). These findings suggest niche microaggregates expanded due to accumulation of beads displaced from flat surfaces.

By contrast, beads located at GFP-expressing AT1 cells were immobile (**Fig. 4E**), indicating that AT1 cell CFTR^null^ expression blocked bead transport. Even within single alveoli lined on opposite walls by TdTomato- and GFP-expressing AT1 cells, only those beads located at TdTomato-expressing AT1 cells were displaced (**Fig. 4G**), ruling out the possibility that bead immobilization at CFTR^null^-expressing AT1 cells resulted from microenvironmental factors. These data provide the first evidence that AT1 cell CFTR function is critical to the convective clearance of particles from alveoli.

Further analysis shows the bead subgroups most immobilized by AT1 cell CFTR^null^ expression were small clusters (**Fig. 4H**) and beads located at flat septa (**Fig. 4I**). AT1 cell CFTR^null^ expression had little effect on bead groups at niches (**Fig. 4I**), where the niche location already restricted bead group movement (**Fig. 4F**). Taking the beads data together, we interpret that AT1 cell CFTR function convectively transports small particle clusters from flat septa toward airways and alveolar niches, efficiently displacing them from sites of AT1 cell contact, collecting them at alveolar corners, and clearing them from alveoli.

### AT1 cell CFTR function clears *S. aureus* from alveoli

Since AT1 cell CFTR function cleared beads from alveoli, we considered it might also clear bacteria. To test this possibility, we intranasally-instilled mice with GFP-expressing *S. aureus* (SA^GFP^), then imaged the SA^GFP^ in alveoli of live, perfused lungs 1-4 h after instillation. We chose this timeframe because our published data show CD11b-expressing neutrophils and monocytes are absent from alveolar airspaces of SA-instilled lungs (*7*), reducing the possibility that innate immune responses would confound SA^GFP^ clearance measures. To avoid fluorescence overlap, we instilled SA^GFP^ in non-fluorescent transgenic mice (AT1-CFTR^null^) that carried *Ager-CreER^T2^* and *Cftr^fl10/fl10^* but not *ROSA^mT/mG^*. Littermate control (LMC) mice carried *Cftr^fl10/fl10^*.

Confocal images of live alveoli of AT1-CFTR^null^ and LMC mice at 1 h after SA^GFP^ instillation show inhaled SA^GFP^ organized against alveolar walls as small clusters of roughly 1-3 bacteria and microaggregates of 4 or more bacteria (**Fig. 5A**). Microaggregates of SA^GFP^ were resistant to alveolar clearance in both groups (**Fig. 5B**) and, thus, unaffected by AT1 cell CFTR^null^ expression. However, AT1 cell CFTR^null^ expression blocked the clearance of small clusters of SA^GFP^ (**Fig. 5B**). These findings show AT1 cell CFTR function clears alveoli of small bacterial clusters. We interpret that loss of AT1 cell CFTR function causes retention of inhaled bacteria against alveolar walls, prolonging bacterial-AT1 cell contact and promoting lung infection.

**Fig. 5.**
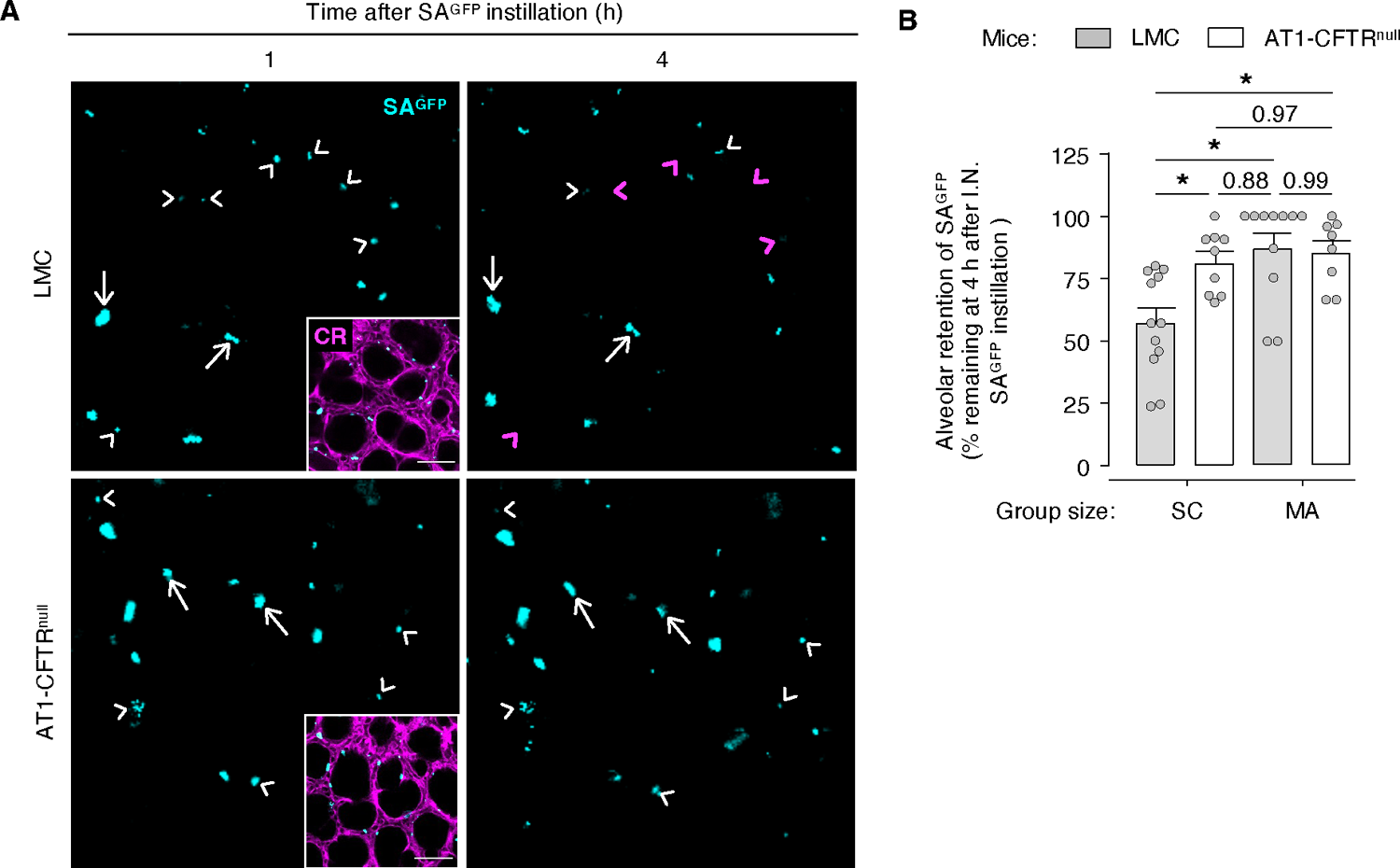
AT1 cell CFTR function clears *S. aureus* from alveoli. (**A-B**) Mice were intranasally-instilled with SA^GFP^, then the lungs were excised for live imaging 10 min later. Confocal images (A) and group data (B) show microaggregates (*arrows*) and small clusters (*arrowheads*) of GFP-expressing *S. aureus* (*SA^GFP^*) in live alveoli of intact, perfused lungs of LMC or AT1-CFTR^null^ mice as indicated. Calcein red-orange (*CR*) dye was added to the perfusate solution to delineate alveolar walls (*inset*). Images were generated in the same alveolar location at 1 and 4 h post-instillation. In the 4 h images, *magenta* arrows and arrowheads point out the initial locations of microaggregates and small clusters that were displaced during the imaging period or were not found in the 4 h images. *White* arrows and arrowheads in the 4 h images indicate microaggregates and small clusters that did not change location. Bar diagrams in B show data from 3 LMC and 3 AT1-CFTR^null^ mice. Circles indicate *n* and each represent one imaging field of at least 30 alveoli; bars: mean ± SEM; *p* as indicated or < 0.05 (*) by ANOVA with post hoc Tukey testing. Scale bar: 50 µm.

## Discussion

Our findings reveal novel and essential roles for AT1 cells in lung homeostasis by showing AT1 cell CFTR function is critical to alveolar liquid secretion and the alveolar clearance of particles and inhaled *S. aureus*. Since AT1 cells comprise over 95% of the lung luminal surface (*20*), we interpret AT1 cell CFTR function is essential to lung health and defense. These findings challenge the widely-held view that the airway is the focus of CFTR function in the lung and suggest that alveoli, particularly AT1 cells, are an important site of the CFTR dysfunction that underlies major lung diseases.

CFTR dysfunction due to genetic causes and smoke exposure underlies the pathogenesis of CF (*29*) and features of COPD (*3, 4, 10, 30–32*) – two diseases that, together, affect nearly 400 million people worldwide (*33, 34*). Although the airway epithelium is considered the site of CFTR dysfunction (*3, 29, 30, 35*), the alveolar epithelium is almost certainly affected on account of its vast surface area (*11, 12*), CFTR expression (*14–19*), liquid secretion function (*6, 13*), and interaction with inhaled particles (*36*). Hence, the alveolar epithelium, and AT1 cells specifically, likely play a major role in CFTR-related disease pathogenesis. Although reports indicating AT1 cells contain little *Cftr* mRNA (*17*) have cast doubt on the significance of AT1 cells in CFTR-related diseases, here we document, by direct observation in live lungs, the physiological importance of AT1 cell CFTR function by showing that AT1 cell CFTR is critical to homeostatic alveolar liquid secretion and alveolar defense.

Our data uncover a seemingly paradoxical relationship between *Cftr* expression and CFTR function in the alveolar epithelium. Human lung AT2 cells outnumber AT1 cells by 2:1 (*37*), and rat AT2 cell *Cftr* mRNA outweighs that of AT1 cells by 5:1 (*17*). These ratios suggest AT1 cell CFTR content is only one-tenth that of AT2 cells, raising the question of how AT1 cell CFTR drives alveolar liquid secretion. One possibility is *Cftr* mRNA quantifications underestimate CFTR function in AT1 cells, since other factors – e.g., CFTR ion channel regulation, CFTR plasma membrane stability, and intracellular Cl^-^ ion content – might favor AT1 cell CFTR function *in vivo.* Another possibility is the 5:1 mRNA ratio overestimates CFTR function in AT2 cells, since immunohistochemical studies (*18*) show considerable AT2 cell CFTR protein is in the cytosol, not at luminal surfaces. Although relationships between *Cftr* expression and CFTR protein function in AT1 and AT2 cells remain unclear, our findings show that AT1 cells, a cell type known for relatively low *Cftr* mRNA content, dominate CFTR-dependent liquid secretion in alveoli.

Our findings establish flat alveolar septa as lung structures critical to host defense, and they raise the possibility that AT1 cell CFTR dysfunction at flat septa underlies lung infections in people with CF and COPD. Our data show CFTR-dependent liquid secretion by AT1 cells defends against particle stabilization in alveoli by convectively displacing small bead clusters from flat alveolar walls, clearing them toward the airways or directing them to niches. The same mechanism defends against alveolar infection, since AT1 cell CFTR function cleared alveoli of small clusters of inhaled SA^GFP^. Our finding that flat septa are physiologically significant sites of CFTR function is supported by histological data that demonstrate CFTR expression on flat surfaces of human alveoli (*18, 38*). Hence, we propose that in healthy lungs, AT1 cell CFTR function at flat septa: (i) displaces inhaled bacteria and particles to minimize their interaction with AT1 cells; (ii) clears bacteria and particles toward airways, where the mucociliary escalator removes them from the lungs; and (iii) gathers bacteria and particles at niches, where AT2 cells might facilitate their immune-mediated elimination via surfactant opsonization (*5*) or fibroblast-endothelial communication (*39*). Loss of these defense mechanisms in lungs with AT1 cell CFTR dysfunction might prolong bacterial and particle contact with AT1 cells and set the stage for lung infection and lung injury.

Our findings suggest therapies intended to promote or restore lung liquid secretion should target AT1 cells. AT1 cell targeting might be achieved by delivering CFTR modulator drugs or expression vectors in preparations that bind AT1 cell surface proteins, like AGER (*24*) and EMP2 (*40*). AT1-targeted inhaled therapies might be designed to deposit in alveoli (*41, 42*), stabilize at flat septa, and account for effects of age, body habitus, lung disease, and alveolar structure on inhalation patterns (reviewed in (*43*)). Whatever their design, AT1-targeted therapies for CFTR-related diseases are likely to augment lung liquid secretion in a critical cell type and thus have an outsized therapeutic effect.

Our study has limitations. First, due to imaging constraints related to the *ROSA^mT/mG^* allele, the dextran we used as an airspace indicator of alveolar liquid secretion was of a different molecular weight (10 kDa) and concentration (10 mg/mL) than the dextran in the perfusate solution (70 kDa, 40 mg/mL). This difference might have generated an osmotic gradient across the alveolar barrier that favored liquid absorption, leading us to underestimate secretion rates. This difference does not invalidate our conclusions, because any effect it had would occur in a cell type-independent manner and bias against our finding secretion differences. Second, mice differ from humans in the airway expression of non-CFTR ion transport proteins (*44, 45*). The extent to which such proteins are expressed in the alveolar epithelium is unclear, and there is little data to indicate whether they contribute to alveolar liquid dynamics in any species. There is no evidence to indicate our mouse lung data do not represent alveolar function in human lungs. Nevertheless, studies in intact, perfused human, pig, or ferret lungs might bolster our findings.

In conclusion, our findings suggest a paradigm shift in our understanding of the role of AT1 cells in lung health and disease. Our data reveal that AT1 cells, long considered to be passive conduits for gas exchange, are dynamic cells that drive homeostatic lung liquid secretion and have a critical role in pathogen clearance and host defense. These findings instruct that loss of AT1 cell CFTR function may underlie lung diseases that affect hundreds of millions of people worldwide, and they suggest that existing therapies that target CFTR in the lung airways overlook the immense therapeutic potential of targeting CFTR in lung alveoli. Going forward, new therapies designed to promote lung liquid secretion for CFTR-related diseases, such as CF and COPD, should consider AT1 cells key therapeutic targets.

## Acknowledgements

General: We thank Dr. Maor Sauler for helpful discussions related to approaches for quantifying *Cftr* in transgenic mice.

## Funding

National Institutes of Health grant R01HL164821 (JH) National Institutes of Health grant R00HL155785 (JK) Cystic Fibrosis Foundation Research Grant 004792G222 (JH) American Lung Association COVID-19 & Emerging Respiratory Viruses Research Award 1031520 (JH) American Lung Association Hastings Innovation Award HIA-1278383 (JK)

## Author contributions

Conceptualization: JH

Investigation: SS, SH, DC, ST, SM, JK, JH Visualization: SS, JH

Funding acquisition: JH, JK Project administration: JH, JK Supervision: JH, JK, CB, AP, AM Writing – original draft: JH

Writing – review & editing: JH, SS, SH, DC, ST, SM, CS, JZ, CB, AP, AM, JK

## Competing interests

Authors declare that they have no competing interests.

## Data and materials availability

All data are available in the manuscript or the supplementary materials.

## Materials and Methods

### Experimental design

The experiments were designed and the manuscript was written according to ARRIVE guidelines. Sample sizes and outcome measures are indicated in figures and legends.

### Fluorophores

We purchased LysoTracker Green (100 nM), AlexaFluor 647-conjugated dextran (AF647; 10 kDa; 10 mg/mL), fluorescein isothiocyanate (FITC)-conjugated dextran (20 kDa; 5 mg/mL), and calcein red-orange AM (10 μM) from ThermoFisher Scientific.

### Reagents

All reagents were freshly constituted for experiments. Tamoxifen was purchased from Sigma-Aldrich, reconstituted at 20 mg/mL in sunflower oil (Sigma), and administered within 30 min of reconstitution. Fluorescent microspheres (1.00 μm, Flash Red, cat. no. FSFR004) were purchased from Bangs Laboratories, Inc.

### Solutions

We purchased Ca^2+^- and Mg^2+^-containing PBS from Corning. Isolated lungs were perfused with HEPES-buffered vehicle solution of pH 7.4 and osmolarity 295 mOsm and containing 150 mM Na^+^, 5 mM K^+^, 1 mM Ca^2+^, 1 mM Mg^2+^, 10 mM glucose, 4% dextran (70 kDa; Molecular Probes), and 1% FBS (Gemini Bio-Products). Fluorophores and reagents microinstilled into alveoli were dissolved or suspended in HEPES-buffered vehicle solution without dextran or FBS.

### Bacterial strain and preparation

*S. aureus* was GFP-expressing USA300 LAC (SA^GFP^). SA^GFP^ were stored at −80°C in 25% glycerol in autoclaved Luria-Bertani (LB) broth media (MP Biomedicals) and propagated on LB-agar plates containing chloramphenicol (10 ug/mL). Plates were refreshed from frozen stock every 2 weeks. Single colonies of SA^GFP^ were propagated in autoclaved LB media containing chloramphenicol (10 μg/mL) in a shaking incubator at 37°C and 200 rpm (New Brunswick Scientific) for 18 h (stationary growth phase), then the bacterial solution was prepared for intranasal instillation by diluting 1.3 mL in 200 uL of PBS containing Ca^2+^ and Mg^2+^.

### Animals

Transgenic mice were on a C57BL/6J background. Strains include: tamoxifen-inducible, *Sftpc-CreER^T2^*(*21*) (Jackson Laboratory strain #028054); tamoxifen-inducible, *Ager-CreER^T2^* (*24*) (Jackson Laboratory strain #032771); *Cftr^fl10^* (*22*) (Case Western Reserve University strain *Cftr^tm1Cwr^*); and *ROSA^mT/mG^* (*23*) (Jackson Laboratory strain #007676). Tail biopsy was performed at 2 weeks of age and sent to Transnetyx for genotyping. All mice were genotyped. To induce *Cre* expression, we gave intraperitoneal injections of 100 μL of tamoxifen solution in sunflower oil at 8 weeks of age. Mice used for experiments were 16-18 weeks old for AWL studies and 28-31 weeks old for beads and bacterial studies. All mice were male except one, which was one of three mice used in the LMC-TdTomato group for AWL experiments.

### Intranasal instillation

SA^GFP^ were prepared for intranasal instillation and instilled within 30 min of bacterial removal from the incubator. Mice were anesthetized with inhaled isoflurane (4%) and intraperitoneal injections of ketamine (up to 1 mg) and xylazine (up to 0.1 mg). Each mouse was instilled with 30 μL of prepared SA^GFP^ solution to deliver 1 x 10^8^ CFU per mouse. Instillation quality was recorded at the time of instillation by the performing investigator and considered acceptable for experiments if no loss of instillate was observed. Mice woke from anesthesia within 3 min of instillation. Within 10 min of intranasal instillation, mice were re-anesthetized with inhaled isoflurane (4%) and intraperitoneal injections of ketamine (up to 100 mg/kg) and xylazine (up to 5 mg/kg) for euthanasia and surgical preparation of isolated, perfused lungs.

### Isolated, perfused lungs

We anesthetized mice with inhaled isoflurane (4%) and intraperitoneal injections of ketamine (up to 100 mg/kg) and xylazine (up to 5 mg/kg), then gave intracardiac injections of heparin (50 units; Mylan) and exsanguinated the mice by cardiac puncture. Using our reported methods (*7, 25*), we cannulated the trachea, pulmonary artery, and left atrium of the heart, then excised the heart, lungs, and cannulas en bloc. Lungs were inflated with room air through the tracheal cannula and perfused through the pulmonary arterial and left atrial cannulas at 0.4-0.6 mL/min with autologous blood diluted in a solution of 4% dextran (70 kDa; Molecular Probes), 1% FBS (Gemini Bio-Products), and HEPES-buffered solution at pH 7.4, osmolality 320 mOsm/kg, and 37°C. We used in-line pressure transducers (ADInstruments) to maintain constant airway pressure 6 cm H_2_O via a continuous positive airway pressure machine (Philips Respironics) and pulmonary artery and left atrial pressures 10 and 3 cm H_2_O, respectively, via a roller pump (Ismatec). The lungs were positioned to enable micropuncture and imaging of the diaphragmatic surface of the right middle lobe, right caudal lobe, or left lung. Portions of the lung surface that were not used for micropuncture and imaging were covered with plastic wrap to prevent desiccation.

### Alveolar microinstillation

We hand-beveled glass micropipettes (Sutter Instruments) to micropuncture single alveoli under bright-field microscopy, as we have done previously (*7, 25*). Micropunctured alveoli were instilled with fluorophores and reagents in solution, resulting in their spread from the micropunctured alveolus to neighboring alveoli. Microinstillations were performed in 1-3 alveoli bordering each imaging field.

### Live lung imaging and analysis

By our established methods (*7, 25*), we viewed alveoli by confocal microscopy (LSM800; Zeiss) with a 20x water immersion objective (NA 1.0; Zeiss) and coverslip. We used bright-field microscopy to randomly select regions of 30-50 alveoli for microinstillation and imaging. All images were acquired as single images using Zen (v.2.6; Zeiss) and recorded as Z-sections. Analyzed images were 4-8 μm below the pleura. Optical thickness was 1 μm for beads imaging and 2 μm for all other experiments, and frame size was 512 x 512 pixels. We established laser, filter, pinhole, and detector settings at the beginning of each imaging experiment to optimize alveolar fluorescence and avoid fluorescence saturation, then maintained the settings for the duration of the experiment. We confirmed absence of bleed-through between fluorescence emission channels. Images were analyzed using ImageJ (NIH; v.2.0.0-rc-69/1.52n). Linear adjustments of brightness and contrast were applied to individual color channels of entire images and equally to all experiment groups. We did not apply downstream processing or averaging.

### Alveolar permeability determination

To determine alveolar barrier properties, we added fluorescein isothiocyanate (FITC)-conjugated dextran (20 kDa; 5 mg/mL) to the intact lung perfusate solution using our established methods (*7, 25*).

### Airspace dextran fluorescence determination

We used our established approach (*7, 25*) to microinstill alveolar airspaces of *ROSA^mT/mG^*-expressing transgenic mice with AF647-conjugated dextran (10 kDa; 10 mg/mL in HEPES-buffered vehicle solution). Provided alveolar barrier function is intact, time-dependent loss of dextran fluorescence indicates dilution by AWL secretion (*6, 7*). To quantify the secretion, we used ImageJ (NIH; v.2.0.0-rc-69/1.52n) software to measure dextran gray levels in alveolar airspaces at 2 and 30 min after dextran microinstillation. To account for heterogeneity of dextran fluorescence changes across imaged alveolar regions, dextran gray levels were quantified as a mean of dextran fluorescence in representative alveoli located at the image top left, top right, bottom left, bottom right, and center. Measurement locations were the same across time points. All experiments were followed by a determination of alveolar barrier permeability using the methods indicated above. We excluded alveolar regions from the dextran analysis if: initial dextran gray levels were less than 20; TdTomato fluorescence intensity increased by more than 10% during the imaging period; or permeability determinations showed leak of microvascular fluid into alveolar airspaces.

### Beads quantifications

We microcentrifuged 1 mL of fluorescent microspheres (“beads”; cat. no. FSFR004; Bangs Laboratories, Inc.) and resuspended them in 20 μL of HEPES-buffered vehicle solution. Beads in solution were microinstilled into alveolar airspaces by alveolar micropuncture under bright field microscopy, then we used confocal microscopy to generate image Z-stacks from 30 min to 3 h post-instillation. *ROSA^mT/mG^* fluorescence delineated alveolar walls. To determine bead group size, we used ImageJ (NIH; v.2.0.0-rc-69/1.52n) software to measure the two-dimensional area of bead groups in single Z-stack images, selecting the Z-stack image in which the bead group was the largest. To define bead group displacement, we used ImageJ to measure the distance from the location of the center of the beads group on the 30 min image to the location of the center of the beads group on the 3 h image. Lateral displacement was defined as the distance (μm) moved between Z-section images that were generated at the same sub-pleural depth; vertical displacement was defined as the distance (μm) moved between Z-section images generated at different depths, with each Z-section image set at 1 μm apart; and total displacement was the sum of the lateral and vertical displacements.

### Bacterial quantification

We added calcein red-orange (10 μM) to the lung perfusate solution for 30 min, then replaced the perfusate with fluorophore-free perfusate solution. Within 1 h of intranasal SA^GFP^ instillation, we identified live alveoli with SA^GFP^ and generated confocal images as Z-sections. Additional Z-sections were generated in the same locations at 4 h post-instillation, then we compared SA^GFP^ fluorescence in the 1 and 4 h post-instillation images at 6 μM below the pleura. We restricted the analysis of SA^GFP^ dynamics to alveoli in which airspace diameters were stable from 1-4 h post-instillation. SA^GFP^ were considered to be retained in alveoli if, on the 4 h post-instillation image, they were visible on the same Z-section and located within 5 μM of their original location on the 1 h post-instillation image. SA^GFP^ that merged with calcein fluorescence were considered to be intracellular and excluded from the analysis.

### Mouse lung epithelial cell preparation and fluorescence-activated cell sorting (FACS) of AT2 cells from mouse lungs

We applied our standard methods (*46, 47*) to isolate AT2 cells from lung homogenate. Single cell suspension of mouse lungs was created as previously described (*48*). Briefly, lungs of euthanized mice were instilled with 1 mL of 15 U/mL Dispase II (ThermoFisher) and incubated with additional 2 mL/mouse of Dispase for 45 minutes at room temperature with agitation. Single cell suspensions were prepared from digested lungs by incubating with DNAse I (10 min at 37°C) and red blood lysis buffer (2 min at room temperature) and filtration through 100 µm, 70 µm, and 40 µm filters. For FACS analysis, single cell preparations were incubated with the following antibodies for 45 min at 4°C in DMEM without phenol red + 2% FBS: Rat anti-mouse CD45 (1:200 dilution, BD Biosciences clone 30-F11), rat anti-mouse CD31 (1:200 dilution; BioLegend clone 390, cat. no. 102434), rat anti-mouse EpCAM (1:500 dilution, BioLegend clone G8.8), and rat anti-mouse integrin β4 (1:75 dilution, BD Biosciences clone 346-11A). Cells were then washed twice with 1X PBS and flash-frozen for RNA isolation. Sorting and subsequent analyses were performed using BD FACS Aria cytometers and FlowJo software.

### qPCR of Cftr RNA

RNA was extracted from cells using PicoPure RNA Isolation Kit (Applied Biosystems cat. no. KIT0204). cDNA was synthesized from total RNA using SuperScript Strand Synthesis System (ThermoFisher cat. no. 18080044). qPCR was performed using SYBR Green (ThermoFisher cat. no. F415L). Relative gene expression levels were defined using the ΔΔCt method. qPCR primers were acquired from Integrated DNA Technologies and are listed in Table S2.

### Statistics

Statistics are indicated in figures and legends. In general, paired comparisons were analyzed using two-tailed *t* tests, and multiple comparisons were made using ANOVA with post hoc testing. We considered statistical significance at *p*<0.05. Data were analyzed and figures were prepared using Microsoft Excel, StatPlus:mac Pro (AnalystSoft, Inc., Build 7.5.0.0/Core v7.6.11), and SigmaPlot (Systat, version 14.5).

### Study approval

The Institutional Animal Care and Use Committee of the Icahn School of Medicine at Mount Sinai approved the animal procedures.

---

**Fig. S1.**
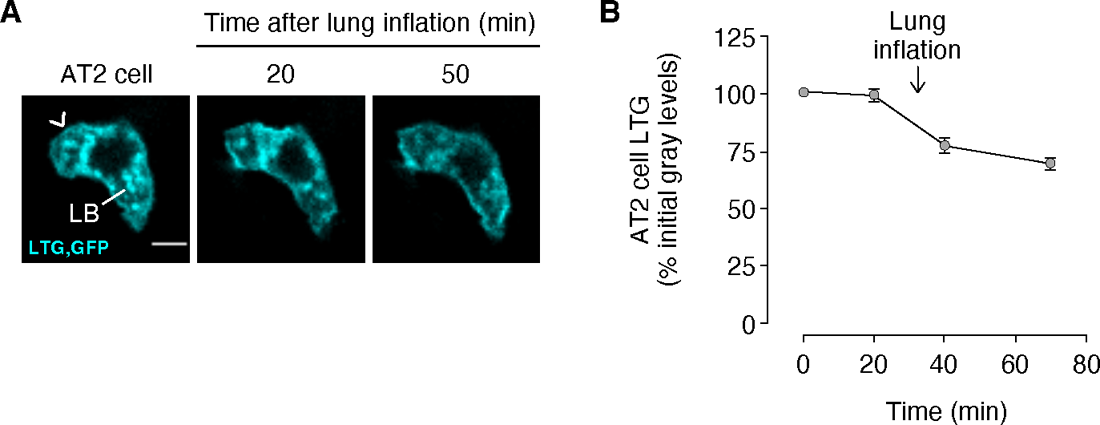
Functional phenotyping of AT2 cells in transgenic mice. (**A-B**) Confocal images (A) show fluorescence of AT2 cell lamellar bodies (LBs) loaded with lysotracker green (LTG) before (*left*) and after (*middle* and *right*) transient lung hyperinflation in an AT2-CFTR^null^-GFP mouse. Plasma membrane GFP fluorescence (*arrowhead*) confirms *Sftpc* expression. Note, LTG fluorescence loss indicates hyperinflation-induced surfactant secretion and affirms AT2 cell identity. Both LTG and GFP are pseudocolored cyan due to fluorescence overlap. In B, circles indicate *n* and each represent one imaging field of at least 30 alveoli in which LTG fluorescence was quantified in at least 3 AT2 cells. Bars: mean ± SEM; *p* values as indicated by two-tailed *t* test. Scale bar: 5 µm.

**Fig. S2.**
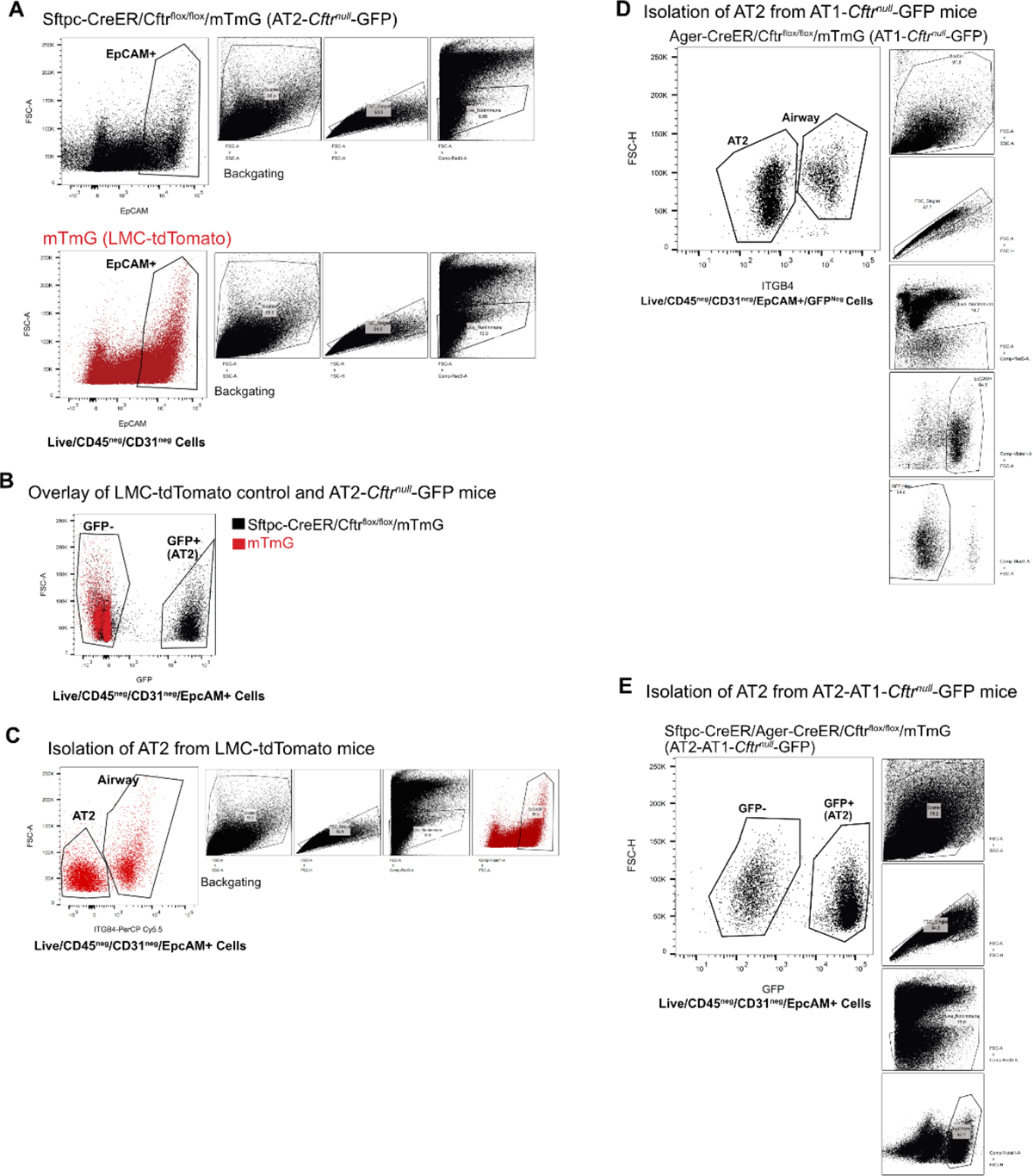
Flow sorting strategy of AT2 cells from lung homogenate. **(A)** Flow plot shows strategy for identifying EpCAM+ cells within Live/CD45^neg^/CD31^neg^ cells prepared from single cell suspensions from tamoxifen-labeled AT2-CFTR^null^-GFP mice (*top panel*) and tamoxifen-labeled LMC-TdTomato mice (*bottom panel*). Backgating shows similar levels of live/non-immune cells. **(B)** Overlay of EpCAM+ cells from AT2-CFTR^null^-GFP and LMC-TdTomato mice against GFP expression shows specificity of tamoxifen-induced GFP expression. GFP+ cells were isolated as AT2 cells from the AT2-CFTR^null^-GFP mouse. **(C)** Strategy of isolating AT2 cells from LMC-TdTomato mice where cells were stained for ITGB4 (an airway epithelial marker). EpCAM+/ITGB4^neg^ were isolated as AT2 cells from LMC-TdTomato mice. **(D)** GFP^neg^/EpCAM+/ITGB4^neg^ were sorted as AT2 cells from AT1-CFTR^null^-GFP mice. Less than 5% of GFP+ cells were observed in backgating, consistent with compromised extraction of AT1 cells using standard lung digestion protocols. **(E)** Isolation of AT2 cells from AT2-AT1-CFTR^null^-GFP mice. GFP+ cells were sorted as AT2 cells from these mice.

**Fig. S3.**
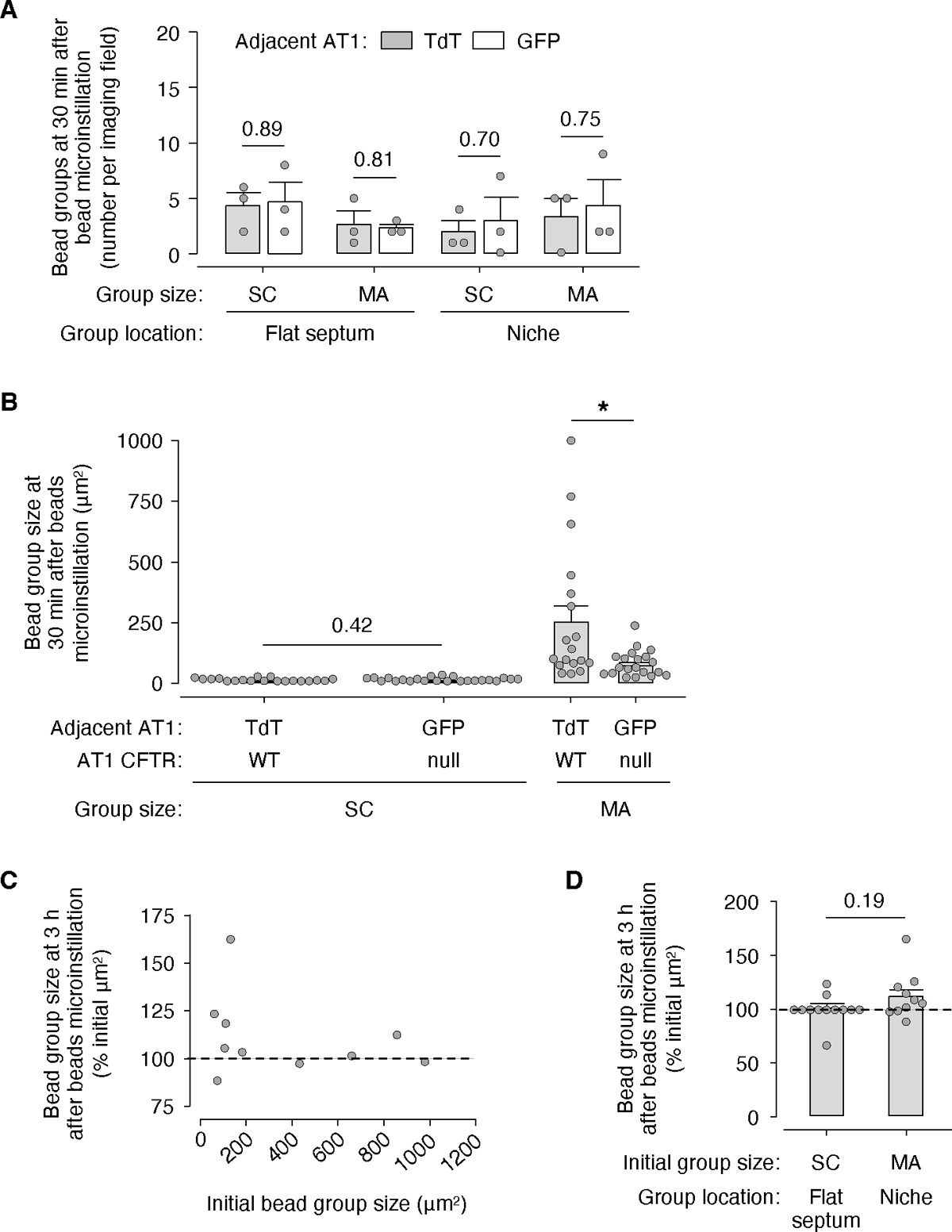
Confocal imaging of fluorescent beads in live alveoli of intact, perfused lungs. (**A-D**) Beads were microinstilled into airspaces of live alveoli of an AT1-CFTR^null^-GFP mouse. We generated confocal images of the beads at 30 min and 3 h post-instillation in 3 imaging fields of at least 30 alveoli and pooled the data to make the plot and bar diagrams. Bead groups were classified as small clusters (*SC*) or microaggregates (*MA*), and their initial locations in alveoli are indicated as either adjacent to TdTomato (*TdT*)- or GFP-expressing AT1 cells. Text in B indicates CFTR expression (CFTR^null^ or WT) signaled by GFP or TdT fluorescence. Circles indicate *n* and each represent means per imaging field (A) or single bead groups (B-D). *Dashed lines* in C-D point out the value that corresponds to no change in bead group size over 3 h after microinstillation. Bars: mean ± SEM; *p* values as indicated or < 0.05 (*) by two-tailed *t* test.

**Table S1.**
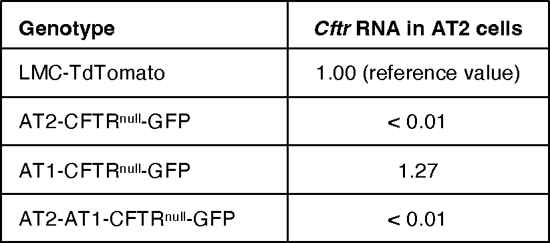
*Cftr* RNA in AT2 cells of transgenic mice. The data show qPCR results by mouse genotype. At least 50,000 AT2 cells were collected per genotype.

| Genotype | *Cftr* RNA in AT2 cells |
| --- | --- |
| LMC-TdTomato | 1.00 (reference value) |
| AT2-CFTR<sup>null</sup>-GFP | < 0.01 |
| AT1-CFTR<sup>null</sup>-GFP | 1.27 |
| AT2-AT1-CFTR<sup>null</sup>-GFP | < 0.01 |

**Table S2.**
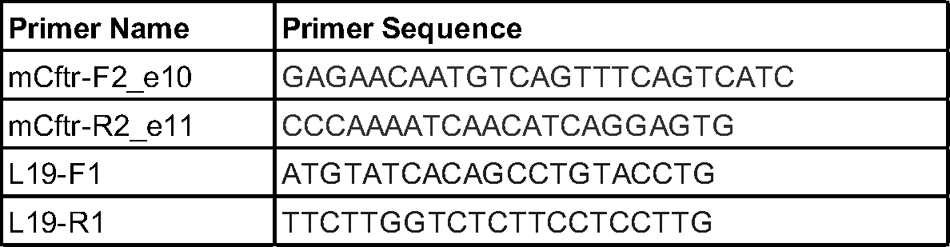
qPCR primers. The indicated primers were used for qPCR of *Cftr* RNA.

## Notes

### Competing Interest Statement

The authors have declared no competing interest.

